# Circadian clock gates retinal regeneration by orchestrating Cxcl12-dependent immune coordination

**DOI:** 10.64898/2026.06.10.731489

**Authors:** Xiangyu Li, Haibo Li, Lianxin Wu, Bowen Lu, Yikun Zhi, Yi Li, Qianyi Lu, Han Wang, Hui Xu

**Affiliations:** Jiangsu Key Laboratory of Tissue Engineering and Neuroregeneration, Key Laboratory of Neuroregeneration of Ministry of Education, Co-innovation Center of Neuroregeneration, Nantong University, Nantong, Jiangsu 226001, China; The Key Laboratory of Pediatric Rare Diseases of the Ministry of Education, University of South China and Department of Cell Biology and Genetics, School of Basic Medical Sciences, Hengyang Medical School, University of South China, Hengyang, Hunan 421001, China; Center for Circadian Clocks, Soochow University, Suzhou 215123, Jiangsu, China, Jiangsu Key Laboratory of Drug Discovery and Translational Research for Brain Diseases, School of Basic Medical Sciences, Suzhou Medical College, Soochow University, Suzhou, Jiangsu 215123, China; Department of Ophthalmology, The First Affiliated Hospital of Soochow University, No.899 Pinghai Road, Suzhou, Jiangsu 215006, China

**Keywords:** retinal regeneration, circadian clock, Müller glia, immune response, Cxcl12

## Abstract

The circadian clock orchestrates tissue homeostasis and repair, yet its role in central nervous system (CNS) regeneration remains largely unexplored. Here, we identify the circadian clock as an essential regulator of adult zebrafish retinal regeneration, a CNS model. Retinal injuries induced at distinct circadian times elicit differential Müller glia (MG) reprogramming and progenitor proliferation, both of which exhibit robust diurnal fluctuations. Disrupting circadian rhythms—via constant darkness (DD) or day-night reversal (DL)—suppresses MG reprogramming, MG-derived progenitor cell (MGPC) formation and neuronal regeneration, concomitant with dampened retinal immune responses. Notably, intravitreal immune stimulation restores MGPC numbers in the retina under DD conditions, linking immune activation to circadian-regulated regeneration. Single-cell RNA-seq reveals that DD disrupts MG reprogramming trajectories while inducing unfolded protein response (UPR) and suppressing ATP/protein synthesis. Moreover, clock disruption alters microglia heterogeneity, and cell-cell communication analysis identifies the Cxcl12-Cxcr4 signaling as a key chemotactic axis between MG/endothelial cell and immune populations. Mechanistically, Bmal1b directly binds E-box motifs in the *cxcl12a/12b* promoters, driving their rhythmic transcription to orchestrate the immune response after injury. Importantly, exogenous Cxcl12a partially restores both immune and regenerative responses in clock-disrupted retinas, establishing the Cxcl12-Cxcr4 axis as a circadian-immune checkpoint for retinal repair. Collectively, our findings reveal a novel paradigm in which the circadian clock gates CNS regeneration through Cxcl12-mediated immune coordination, offering potential therapeutic insights for CNS repair in mammals.

## 1. INTRODUCTION

The circadian clock is an evolutionarily conserved timekeeping system that orchestrates tissue homeostasis and repair across organisms^1,2^. While circadian regulation of injury responses and stem cell behaviors has been demonstrated in non-neuronal tissues such as the skin and liver^3,4^, its role in nervous system repair remains underexplored. In the peripheral nervous system (PNS), circadian modulation of axonal regeneration has yielded conflicting results^5,6^. Whether the circadian clock regulates the regeneration in the central nervous system (CNS) following injury is unknown.

Unlike mammals, teleost fish such as zebrafish possess a remarkable capacity to regenerate damaged CNS tissues, including the retina^7,8^. Following retinal injury, zebrafish Müller glia (MG) undergo reprogramming and asymmetric division to generate MG-derived progenitor cells (MGPCs), which further proliferate and differentiate into retinal neurons to repair the retina ^9,10^. Since the mammalian CNS exhibits extremely limited regenerative capacities after injury, the zebrafish retina serves as a powerful model for investigating how the circadian clock interacts with stem cell plasticity and neuronal regeneration in the vertebrate CNS.

In addition to numerous intrinsic reprogramming factors, extrinsic signals from inflammatory and immune cells are known to prime MG for reprogramming and proliferation in the injured zebrafish retina^11–16^. Given the established roles of circadian clock in immune regulation^17–19^, we hypothesized that it may govern retinal regeneration via immune modulation. Two key questions arise: (1) Are the immune and regenerative responses in zebrafish retina under circadian control? (2) Does the circadian clock gate retinal regeneration via immune modulation? If so, what are the molecular mechanisms linking the clock-dependent regeneration to immune signaling?

Here, we investigate how circadian disruption alters the regenerative process in the injured zebrafish retinas, focusing on immune modulation as a potential gating factor. Using photoperiod manipulation, we uncover that the circadian clock gates retinal repair through Cxcl12-dependent immune coordination. Our work not only reveals a novel circadian-immune crosstalk in CNS regeneration, but also provides a conceptual framework for exploring time-dependent therapies in non-regenerative systems, such as the mammalian retina.

## 2. Materials and Methods

### 2.1 Animals and Eye Injury

All zebrafish used in this study were treated following the Guidelines for Animal Use and Animal Care at Nantong University. Both sexes of adult Wild-type (AB), *Tg(gfap:GFP)*^20^ and *Tg(mpx:GFP)*^21^ transgenic zebrafish at the age of 6 to 8 months were maintained in an automatic breeding system at 28 °C on LD (14 hr light: 10 hr dark), DD (24 hr dark), DL (14 hr dark:10 hr light) or LL (24 hr light) cycles. All of the light manipulations started from 3 days prior injury till 2-4 days post injury (dpi). Methods for introducing mechanical retinal injury via needle-poke have been described in our previous study^11^. For analysis of rhythmic gene expression or cell numbers, retinas were injured at 0:00 am (ZT 16) of 0 d. Retinas in other experiments were injured at 9:00 am (ZT 1) of 0 d.

### 2.2 Immunofluorescence and Lineage Tracing

Tissue preparation, cryosection, and immunofluorescence staining were performed as described previously^22^. The following primary antibodies were used: mouse anti-Zpr1 (1:200, Cat# ab174435, Abcam, Cambridge, MA, USA); and mouse anti-Hu-antigen C/D (HuC/D; 1:500, Cat# A-21271, Thermo Fisher Scientific, Waltham, MA, USA). 1 mg of Dylight 594-labeled isolectin B4 (IB4, Vector laboratories, Inc., Burlingame, CA, USA) was intravitreally injected 24 hours before sacrifice to label retinal microglia. To label proliferating cells, 5-Ethynyl-2’-deoxyuridine (EdU) staining was performed using a BeyoClickTM EdU-488 Cell Proliferation Detection Kit (Cat# C0071L, Beyotime, Shanghai, China). For MGPC lineage tracing, fish received intraperitoneal EdU injection (20 μl of 20 mM, Cat# HY-118411, MedChemExpress LLC, Monmouth Junction, NJ, USA) at 4 dpi before sacrificed at 30 dpi to identify MGPC progeny in the retina.

### 2.3 Quantitative PCR (qPCR)

Fish retinas were collected and total RNA was isolated using the TRIzol reagent (Thermo Fisher Scientific, Waltham, MA). qPCR was performed in triplicate using methods described previously^22^ and *n* ≥ 3. Relative mRNA levels were calculated using the ΔΔCt method^23^ and normalized to the level of *rpl13*. qPCR primers used in the study were listed in Supplementary **Table 1**.

### 2.4 Single-cell RNA sequencing (scRNA-seq) and analysis

Detailed methods for preparing zebrafish retinal samples (0 d, 1 dpi, 2 dpi, and 4 dpi) and scRNA-seq have been described in our previous study^15^. Briefly, six retinas per timepoint were mixed and digested in 1000u/ml papain solution (Solarbio Life Science, Beijing, China) for 10 min at room temperature. Digested cells were further washed, filtered and resuspended in cold 2% FBS (Thermo Fisher Scientific, Waltham, MA, USA). Suspended single-cell solutions were examined for cell vitality (> 80%), adjusted to ∼1000 cells/ml, and loaded onto the 10x Chromium Chip (10x Genomics) for cDNA amplification and library construction. Libraries were sequenced by LC-Bio Technology (Hangzhou, China) on an Illumina NovaSeq 6000 sequencing system.

Sequencing results were analyzed using the OmicStudio tools created by LC-Bio Technology (https://www.omicstudio.cn/cell). Readings from Illumina sequencing were converted to FASTQ format and aligned to zebrafish GRCz11_98 genome using the CellRanger software (https://support.10xgenomics.com/single-cell-gene-expression/software/pipeli <u>nes/latest/what-is-cell-ranger</u>, version 7.0.0). Low quality cells were filtered and cell clusters were projected into 2D space using t-Distributed Stochastic Neighbor Embedding (t-SNE) or Uniform Manifold Approximation and Projection (UMAP). A total of 109,439 retinal cells were counted and the median number of genes per cell was 757-925. Retinal cell types were identified based on known markers^24^. To perform the Gene Ontology (GO) or Kyoto Encyclopedia of Genes and Genomes (KEGG) enrichment analysis, hypergeometric testing was conducted on the differential genes of each cluster obtained from Findallmarker analysis compared to other clusters. Gene Set Enrichment Analysis (GSEA) was performed to identify biological pathways associated with specific cell clusters or treatment condition using the OmicStudio tools. Pseudotime analysis was performed using the Monocle3 software (https://cole-trapnell-lab.github.io/monocle3/). Cell-cell communication analysis was performed using the CellPhoneDB database^25^, where zebrafish gene names were converted to corresponding human homolog genes.

### 2.5 Intravitreous Cxcl12a Injection and Drug Treatment

Recombinant zebrafish Cxcl12a (SDF-1α) protein (Cat# abx178492, Abbexa, Cambridge, UK) was dissolved as 1 mg/ml stocks in PBS. 1 μl of 5 ng/μl Cxcl12a was injected intravitreally through the front of the eye at the time of retinal injury. AMD3100 (Cat#HY-10046, MedChemExpress LLC, Monmouth Junction, NJ, USA) was prepared as a 90 mM stock in ethanol. Fish received a daily intravitreal injection of 1 μl of 1mM AMD3100 until harvest.

The methods for immune manipulation in zebrafish retinas have been described previously^11^. To enhance retinal immune response, fish received intravitreous injection of zymosan A (Zym, 1 μl of 10 mg/ml, Sigma-Aldrich, St. Louis, MO, USA) at the time of retinal injury.

### 2.6 Cell Culture and Luciferase Reporter Assays

Cell culture and luciferase reporter assays were performed as previously described. 2-kb 5’fragments of the *cxcl12a* and *cxcl12b* promoter were amplified with PCR and sub-cloned into a luciferase reporter vector PGL4.17-luc, respectively. The results of prediction by Jaspar^26^ (https://jaspar.genereg.net) showed that the *cxcl12a* promoter contains one E-box (5’-TCACGTG-3’) and the *cxcl12b* promoter contains one E-box (5’-TCACGTG-3’). Deletion of these regulatory elements was done with the Mut Express II Fast Mutagenesis Kit (Cat# C214, Vazyme). PcDNA3.1 plasmids of full-length cDNAs of Clock1a, Bmal1b and Per2 were reported previously^27^, Human Embryonic Kidney (HEK) 293T cells were co-transfected by Lipofectamine 2000 (Cat#11668027, Invitrogen). 24 hours later, cells were collected and luciferase activities were measured using Dual-Report assay system (Cat# E1980, Promega). For each luciferase assays, at least three independent experiments were conducted. The primers used are listed in Supplementary **Table 2**.

### 2.7 Chromatin immunoprecipitation (ChIP) assays

ChIP assays were conducted as previously described^28,29^. Anti-Bmal1b antibody (Bmal1b protein sequence: PSSQLTQSPESDR)^27,28^ and ChIP assay kit (Cat#4000321, Merck) were used to obtain DNA from 5 adult WT brains that interact with Bmal1b. Finally, the ChIP results were analyzed using PCR.

### 2.8 Microscopy and cell counting

Fluorescence images of retinal sections were acquired using a Zeiss Imager M2 microscope (Carl Zeiss AG, Oberkochen, Germany) with either a 10x or 20x objective. Cell counting in each of the four injured retinal region was performed using the Cell Counter plugin in ImageJ Software.

### 2.9 Statistics

The experimental data were analyzed using GraphPad Prism 9.0. For single comparison, a Student’s *t*-test (2 tails, unpaired) was used to compare the experimental group with the control. For multiple comparisons, one-way or two-way analysis of variance (ANOVA) was performed, followed by a Dunnett’s multiple comparison test. Circadian rhythmicity of the data was evaluated using nonlinear regression (curve-fit) analyses with a single-component cosine-wave model. The significance of the rhythm was determined by the zero-amplitude test. This approach is well-established for circadian rhythm analysis and is particularly suitable for post-injury time series where oscillation amplitude may vary across cycles^30,31^. For simplicity, only the significant differences between the peak and trough on each day are marked in the figures. The difference was considered significant at \**p* < 0.05, \*\**p* < 0.01; \*\*\**p* < 0.001. All experiments except the scRNA-seq were repeated at least three times, and results are presented as mean±SEM.

## 3. RESULTS

### 3.1 Zebrafish retina exhibits diurnal fluctuations in regenerative responses following stab injuries

To investigate the potential role of circadian rhythm in retinal regeneration, we first assessed whether the regenerative response varied with the time of injury (day or night). For this purpose, zebrafish retinas were stabbed at either 15:00 (ZT7, day) or 3:00 (ZT19, night) of the first day (0 d), and retinal samples were collected at 48 hr (2 d) and 72 hr (3 d) post-injury, respectively (Figure 1A). qPCR analysis revealed that the retinal expression of key regeneration-associated genes (RAGs), such as *ascl1a, lin28a*, and *cdk1*^11,32^, was significantly lower when the injury was performed during the day compared to that at night (Figure 1B). EdU immunofluorescence revealed a reduced number of progenitor cells in retinas injured at the day compared to those at night (Figure 1C,D). These results indicated a stronger regenerative response when the retinal injury was performed at night rather than during the day.

**Figure 1.**
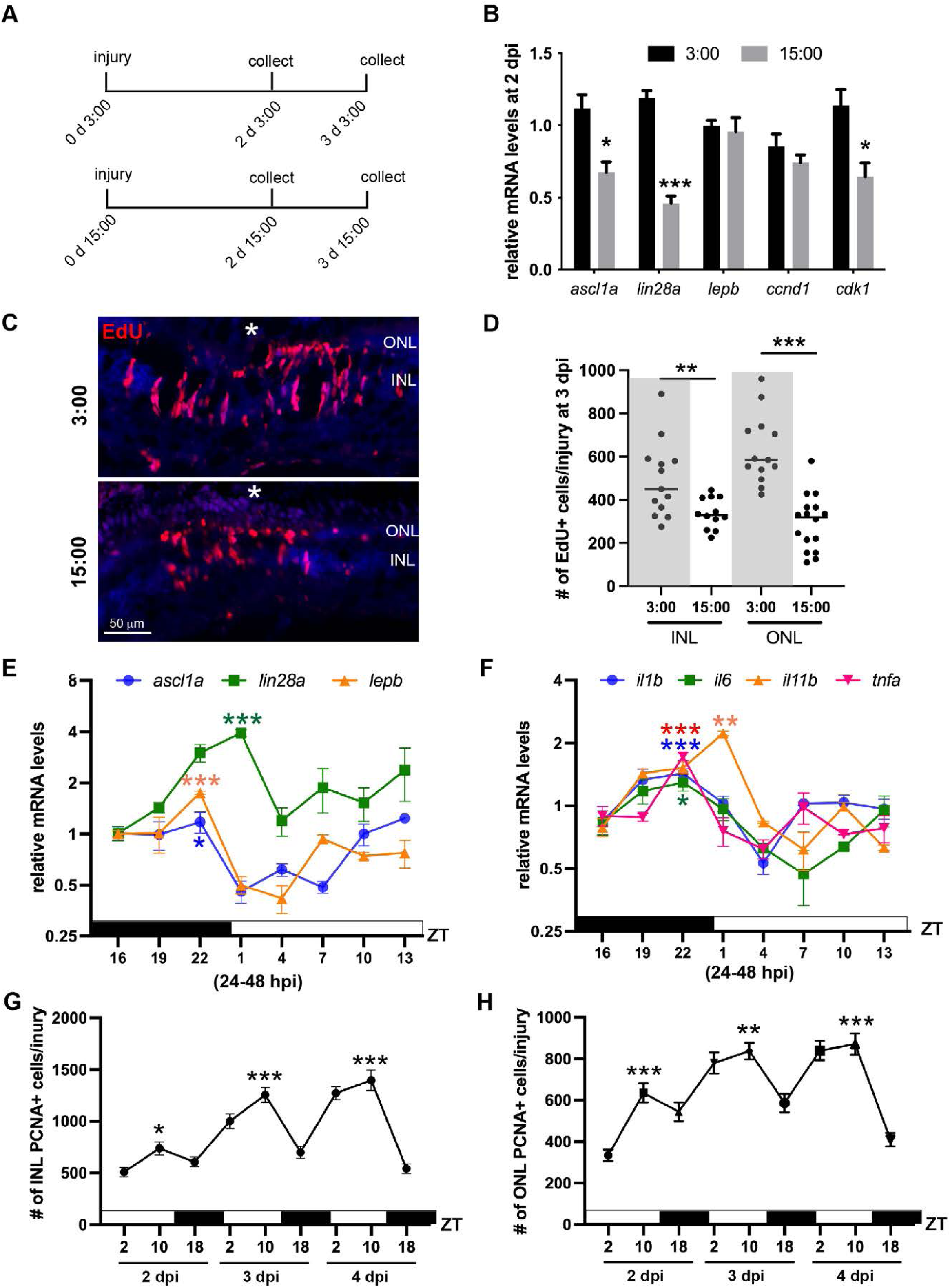
Diurnal fluctuation of the regenerative responses in stab-injured zebrafish retinas. (A) Experimental timeline. Retinas were injured at night (3:00, ZT19) or during the day (15:00, ZT7), respectively. (B) qPCR analysis of regeneration-associated gene (RAG) expression in retinas injured either during the day or at night. (C) EdU immunofluorescence staining showing proliferating cells in the retinas at 3 days post-injury (dpi). White asterisks mark the needle-poke injury sites. (D) Quantification of EdU^+^ cells per injury in (C). (E,F) qPCR analysis showing diurnal fluctuations in the expression of RAGs (E) and inflammatory cytokines (F) in the retina. (G,H) Quantification of EdU^+^ cells per injury across different retinal layers from 2 to 4 dpi. The significant differences between peak and trough time points are indicated in (E-H). *, *p* < 0.05; **, *p* < 0.01; ***, *p* <0.001. ONL, outer nuclear layer; INL, inner nuclear layer.

We next investigated whether the regenerative response exhibited diurnal fluctuations when the retinal injury was performed at a single timepoint (ZT16, corresponding to 0:00). qPCR analysis revealed significant diurnal variations of RAGs and inflammatory cytokines in the retina during 24-48 hours post-injury (hpi) (zero amplitude F-test, *p* < 0.05). These genes exhibited a similar expression pattern, in which their levels gradually increased at night, peaked at around ZT22, and then rapidly decreased to the lowest level at around ZT4, followed by a gradual return to baseline levels (Figure 1E,F). The observed higher expression levels of RAGs and inflammatory cytokines at night were consistent with the results in Figure 1A-D, suggesting that the regenerative and inflammatory responses were stronger during the rest phase (night). EdU fluorescence staining revealed significant rhythmic oscillations (zero amplitude F-test, *p* < 0.0001) in the number of proliferating MGPCs (inner nuclear layer, INL) and rod progenitors (outer nuclear layer, ONL) between 2-4 days post-injury (dpi), in which progenitor proliferation peaked during the day (18:00, ZT10), in contrast with the nocturnal peaks of RAGs expression (Figure 1G,H).Together, these findings demonstrated a diurnal fluctuation of the regenerative response in stab-injured zebrafish retinas.

### 3.2 Circadian clock disruption suppresses retinal regenerative responses

To explore the circadian role in retinal regeneration, light manipulations by constant darkness (DD) or day-night reversal (DL) treatment were used to disrupt the circadian clock starting 3 days prior injury, and their regenerative responses were compared to the control (LD). qPCR analysis demonstrated that under LD conditions, the expression of core clock genes, including *bmal1a*, *clock1a*, *per1b* and *cry1a*, exhibited significant 24-hour rhythmic oscillations from 0-48 hpi in the retina (zero-amplitude F-test, *p* < 0.05, Figure 2A-D). Specifically, the expression levels of *bmal1a* and *clock1a* peaked at ZT4 (day, active phase) and reached a trough around ZT22 (night, rest phase) (Figure 2A,B). In contrast, negative clock regulators such as *per1b* and *cry1a* were expressed in an antiphasic manner to *bmal1a* and *clock1a* (Figure 2C,D). Importantly, both DD and DL treatment significantly reduced the amplitude and mesor (baseline expression levels) of core clock genes in the retina during this period, with no significant phase shift detected (Figure 2A-D).These results confirmed the effectiveness of light manipulation in disrupting the circadian clock.

**Figure 2.**
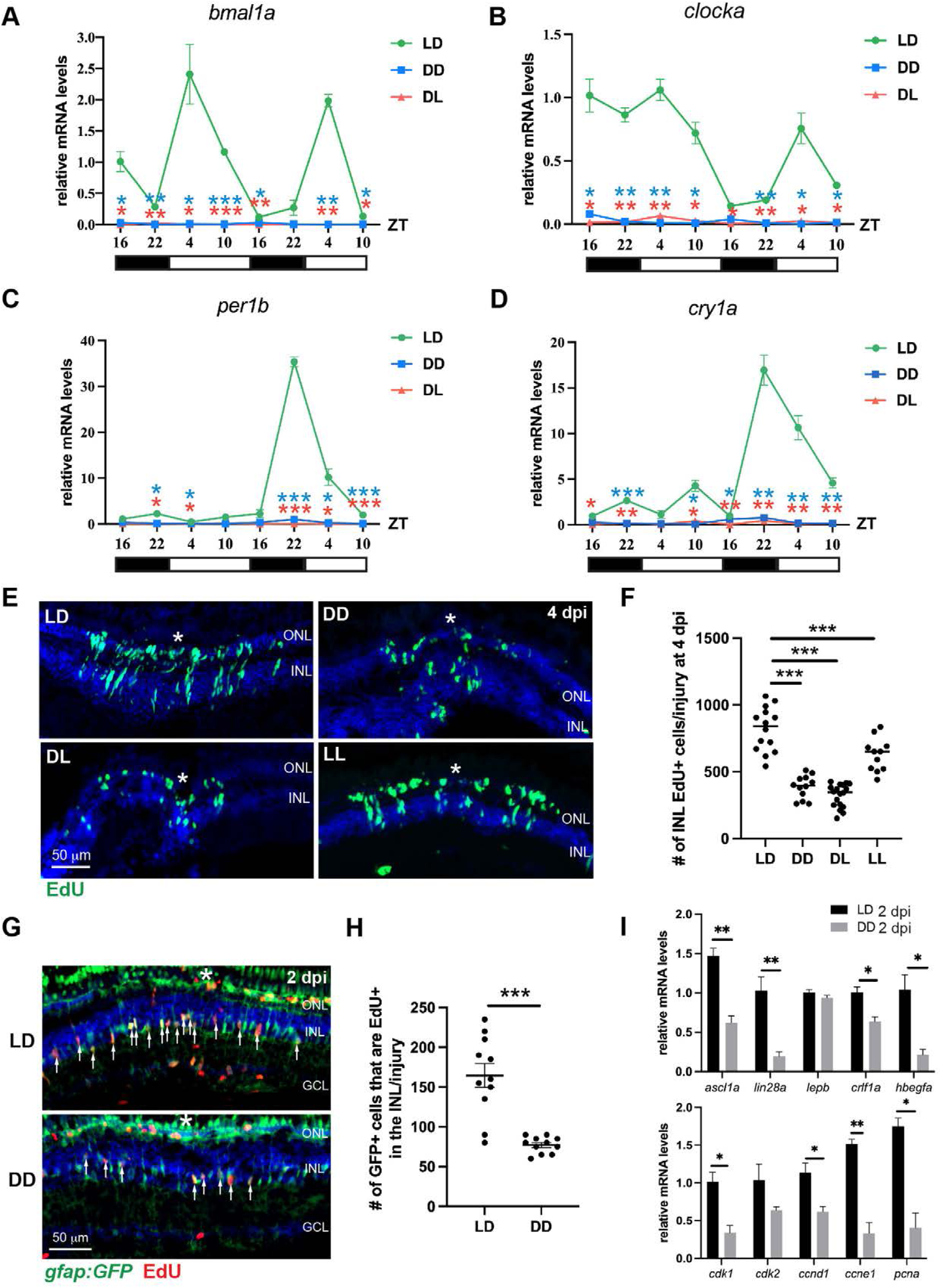
Disrupting the clock suppresses the retinal regenerative responses. (A-D) qPCR analysis of the retinal expression of core clock genes *bmal1a*, *clocka*, *per1b* and *cry1a* from 0-48 hpi under LD, DD or DL conditions. Gene expression levels in DD or DL retinas are compared to those in LD controls. (E) EdU immunofluorescence showing proliferating cells in the INL at 4 dpi (MGPCs). (F) Quantification of EdU^+^ cell number per injury in (E). (G) EdU staining showing proliferating MG (EdU^+^ GFP^+^) in the *Tg(gfap:GFP)* transgenic line. (H) Quantification of proliferating MG in (G). (I) qPCR showing the expression of genes involved in MG reprogramming and cell cycle progression in LD or DD retinas. White asterisks indicate the needle-poke injury sites. *, *p* < 0.05; **, *p* < 0.01; ***, *p* <0.001. ONL, outer nuclear layer; INL, inner nuclear layers; GCL, ganglion cell layer.

We then assessed the impact of circadian clock disruption upon the formation of MGPC clusters at 4 dpi. EdU immunostaining showed that DD-treated retinas exhibited significantly lower number of MGPCs (INL EdU+ cells) compared to LD control at 4 dpi (Figure 2E,F). Since light itself may directly activate the regenerative responses, the observed phenotype in the DD group may be due to lack of light exposure, rather than a disrupted circadian rhythm. To exclude this possibility, we further examined the MGPC number in DL- or constant light (LL)-treated retinas, in which they received a similar (DL) or even larger amount of light exposure (LL) compared to the LD control. Importantly, both DL- and LL-treated retinas exhibited significantly lower number of MGPCs compared to LD control (Figure 2E,F), suggesting the observed regenerative deficiency in the DD group was not due to absence of light exposure. We next investigated the effect of circadian disruption on MG proliferation, which occurred at 2 dpi^14,33^. For this purpose, a transgenic zebrafish line *Tg(gfap:GFP)* was used to label MG in the zebrafish retinas, and EdU immunostaining showed that the number of proliferating MG (GFP^+^/EdU^+^) in DD-treated retinas was significantly lower than that of the LD control (Figure 2G,H). Furthermore, qPCR analysis showed that DD treatment significantly reduced the expression of genes necessary for MG reprogramming^32,34–36^ (Figure 2I, upper panel) and cell-cycle-related genes (Figure 2I, lower panel) in the retina. Together, these results indicated that an undisturbed circadian clock was required for MG reprograming/proliferation and the formation of MGPC clusters in injured zebrafish retinas.

### 3.3 Interference with the circadian clock irreversibly impairs the regeneration of retinal neurons

Since disruption of the circadian clock suppresses MG proliferation and MGPC formation at 2-4 dpi, we asked whether this effect is reversible once the fish were returned to normal light-dark cycles. For this purpose, fish under LD or DD conditions received intraperitoneal EdU injection at 4 dpi to label the proliferating MGPCs, and the DD group was then returned to normal LD cycles after 4 dpi. The progeny cells of MGPCs were lineage-traced till 7 and 30 dpi. EdU immunofluorescence showed that DD-treated retinas exhibited a significantly lower number of MGPC progeny cells (EdU+ cells) than the LD group at 7 and 30 dpi (Figure 3A,B,D,E), suggesting that the effect of circadian disruption on the MGPC population was irreversible. No significant difference in MGPC’s distribution among the three nuclear layers was found between the LD and DD groups (Figure 3C,F), suggesting a dispensable role of circadian clock in MGPC’s cell fate decision. Immunofluorescence staining of the cell type markers of major retinal neurons showed that DD-treatment significantly reduced the number of regenerated retinal neurons at 30 dpi, including photoreceptors (Zpr1^+^ EdU^+^ in ONL), amacrine cells (HuC/D^+^ EdU^+^ in INL), and retinal ganglion cells (RGCs, HuC/D^+^ EdU^+^ in GCL) (Figure 3 G-I). Together, these results revealed an essential role of circadian clock in the regeneration of retinal neurons in adult zebrafish retinas.

**Figure 3.**
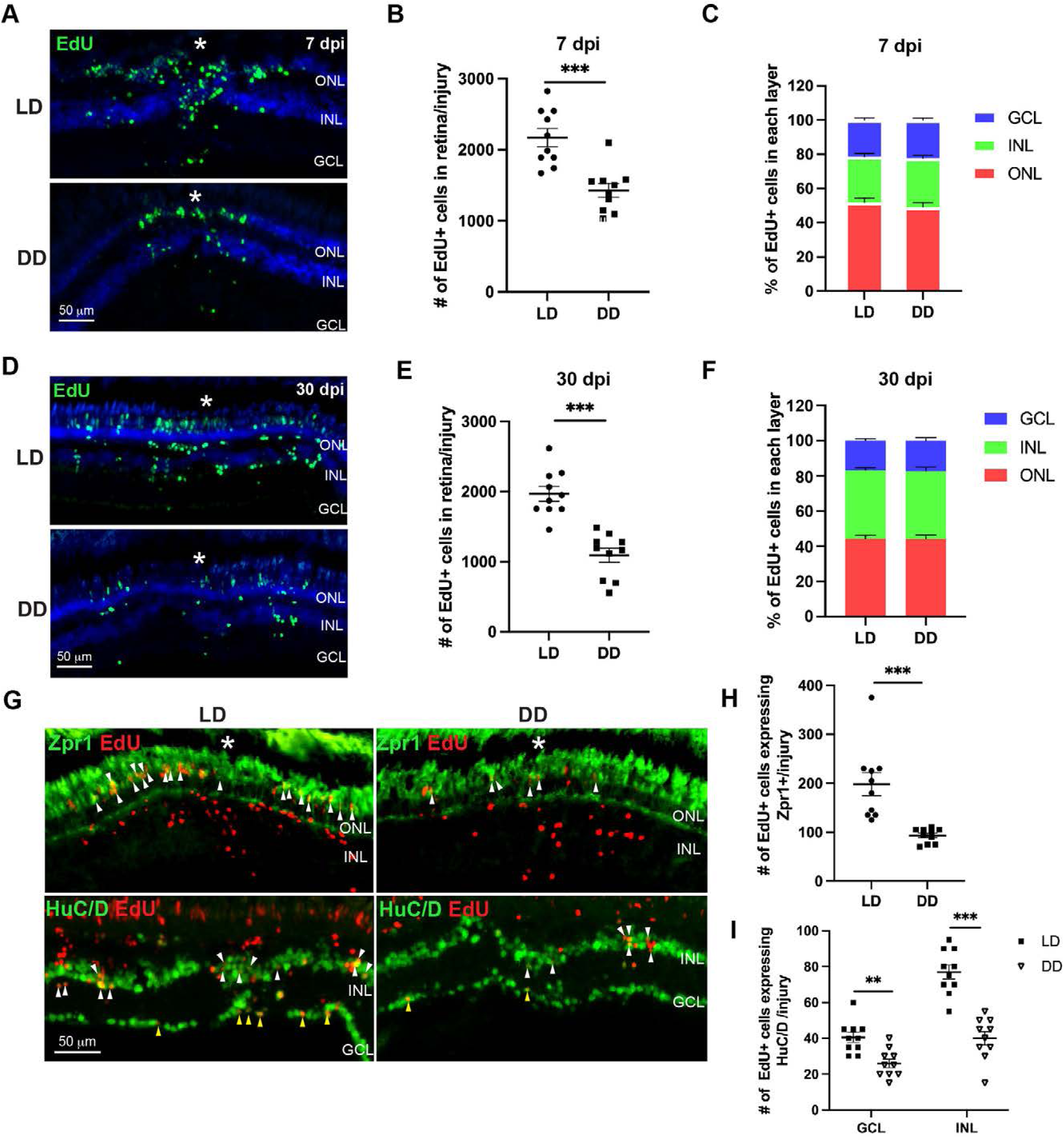
Circadian disruption impairs retinal neuron regeneration. (A) EdU lineage tracing showing MGPCs in the retina at 7 dpi under LD or DD conditions. (B,C) Quantification of the total number (B) and spatial distribution (C) of EdU^+^ cells in the injured retina in (A). (D) EdU lineage tracing showing MGPC progeny in the injured region at 30 dpi under LD or DD conditions. (E,F) Quantification of the total number (E) and distribution (F) of EdU^+^ cells in the injured retina in (D). (G) Immunofluorescence staining for EdU and cell-type-specific markers identifying regenerated retinal neurons at 30 dpi. (H) Quantification of regenerated photoreceptors in (G); (I) Quantification of regenerated amacrine cells (in the INL) and RGCs (in the GCL) in (G). White asterisks mark the needle-poke injury sites. *, *p* < 0.05; **, *p* < 0.01; ***, *p* <0.001. ONL, outer nuclear layer; INL, inner nuclear layers; GCL, ganglion cell layer.

### 3.4 The innate immune response is under circadian clock’s control in injured retinas

The circadian clock plays a crucial role in regulating the reactivity of innate immune cells such as microglia/macrophage and neutrophils^19^. Among these cells, microglia are known to be essential for MG reprogramming and proliferation in zebrafish^11^. To investigate whether the innate immune response was under circadian control in injured zebrafish retinas, we monitored the accumulation of microglia at the injury site every 3 hours between 24 and 48 hpi under LD conditions. Fluorescence microscopy revealed a diurnal fluctuation in the number of IB4^+^ microglia within the injury region: the microglia counts gradually increased during the night, peaked shortly before light onset, and subsequently declined throughout the day (Figure 4A,B). This suggests that microglia reactivity may be regulated by the circadian clock. We next examined the impact of circadian disruption on the innate immune responses following retinal injury. Our results showed that both DD- and DL-treatment significantly reduced the accumulation of IB4^+^ microglia at 1 and 2 dpi, as well as GFP^+^ neutrophils at 3 and 6 hpi in the injured region (Figure 4C-F, Figure S1), suggesting that the chemotaxis of these immune cells to the injury region was under circadian control. DD-treatment also significantly reduced the retinal expression of inflammatory cytokines such as TNFα (*tnfb*), IL-1β (*il1b*), IL-6 (*il6*) and IL-11α (*il11a*) at 2 dpi as shown by the qPCR analysis (Figure 4G). Importantly, intravitreal injection of zymosan A (Zym) to enhance retinal immune responses was able to rescue the deficiency in MGPC formation in DD conditions, as demonstrated by the EdU immunostaining at 4 dpi (Figure 4 H,I), suggesting that circadian clock regulated retinal regeneration via modulating the immune responses. Together, these findings indicated that the innate immune response was controlled by the circadian clock in stab-injured zebrafish retinas.

**Figure 4.**
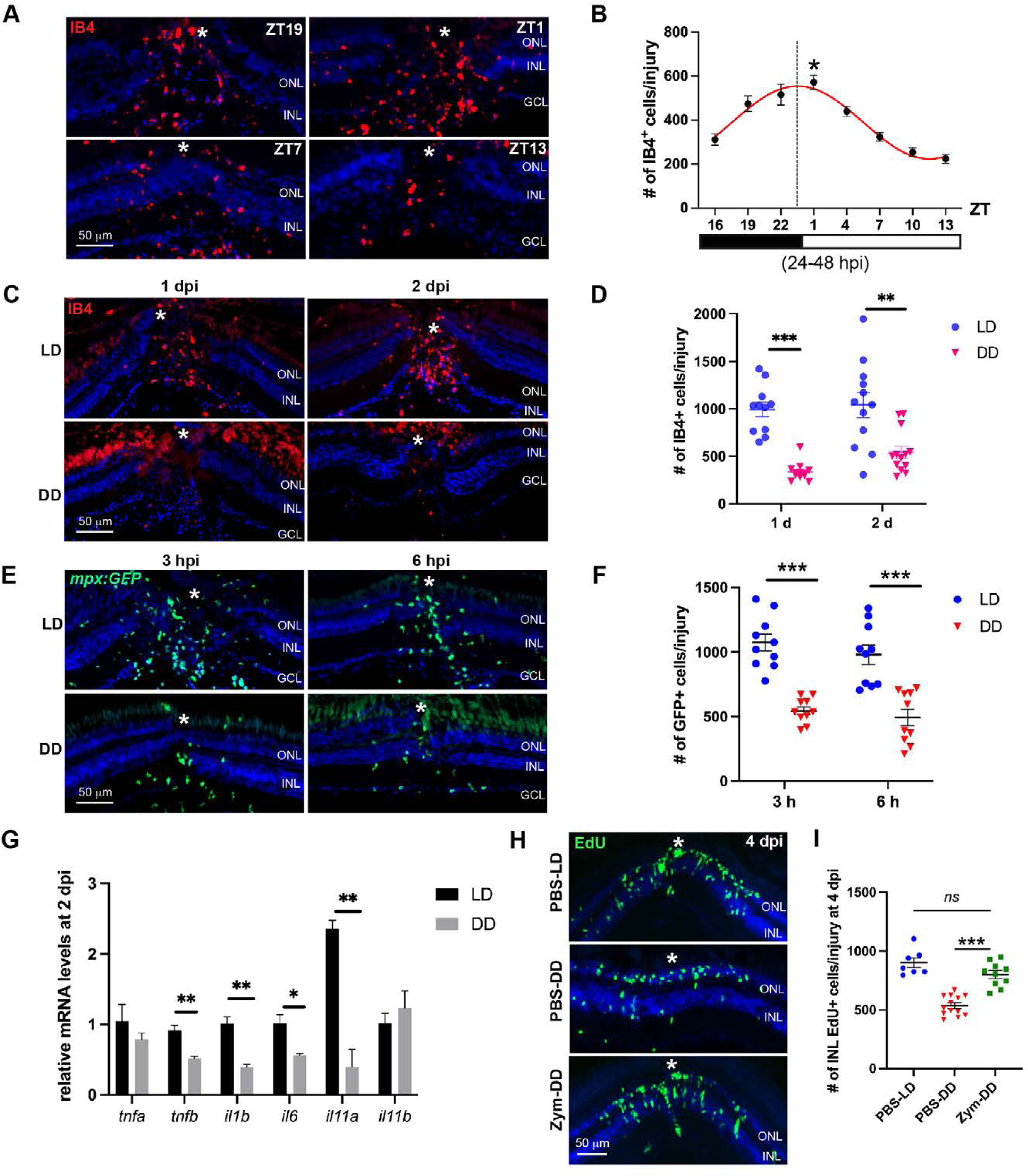
Circadian disruption suppresses the innate immune responses in the injured retina. (A) IB4 immunofluorescence showing microglia accumulation at the indicated time points from 24 to 48 hpi in the injured retina. (B) Diurnal fluctuation of microglia counts at the injury site in (A). Zero-amplitude F-test, *p* <0.0001. The best-fit curve is shown in red, and the peak time is indicated by the dashed line. (C) IB4 immunofluorescence showing microglia at 1 and 2 dpi in the retinas under LD or DD conditions. (D) Quantification of microglia per injury in (C). (E) Distribution of neutrophils in the retina of the *Tg(mpx:GFP)* transgenic line. (F) Quantification of neutrophils per injury in (E). (G) qPCR analysis showing the retinal expression of inflammatory cytokines under LD or DD conditions. (H) EdU immunostaining showing MGPCs in the retina under indicated conditions at 4 dpi. (I) Quantification of MGPCs per injury in (H). White asterisks mark the needle-poke injury sites. *ns*, not significant; *, *p* < 0.05; **, *p* < 0.01; ***, *p* <0.001. ONL, outer nuclear layer; INL, inner nuclear layers; GCL, ganglion cell layer.

### 3.5 scRNA-seq reveals altered MG dynamics and metabolic-stress signatures in DD-treated retinas

To assess the effects of circadian disruption on immune and regenerative responses, we performed scRNA-seq on zebrafish retinas (0-4 d) under LD or DD conditions. A total of 109,439 cells were counted and separated into 34 clusters, and retinal cell types were identified and merged using known markers (Figure S2). Based on known MG status markers^15,24^, we identified six MG subtypes: Resting (Res-MG), Reactive (Rea-MG), Activated (Act-MG), Immunoregulatory (IR-MG), Proliferating (Pro-MG), and Post-mitotic (PM-MG) (Figure 5A-C). Among them, the Res-MG, Act-MG and IR-MG subgroups have been described previously^15^. The Rea-MG, enriched in injured LD retinas, highly expressed gliosis marker *gfap* and mitochondrial ATP synthesis genes (Figure 5C,D). Compared to Act-MG, Rea-MG lacked regeneration-associated gene (RAG) expression such as *lepb*, *crlf1a* and *yap1* (Figure 5C,D), suggesting that it was either a subgroup of reactive but non-regenerative MG, or an intermediate subtype between Res-MG and Act-MG.

**Figure 5.**
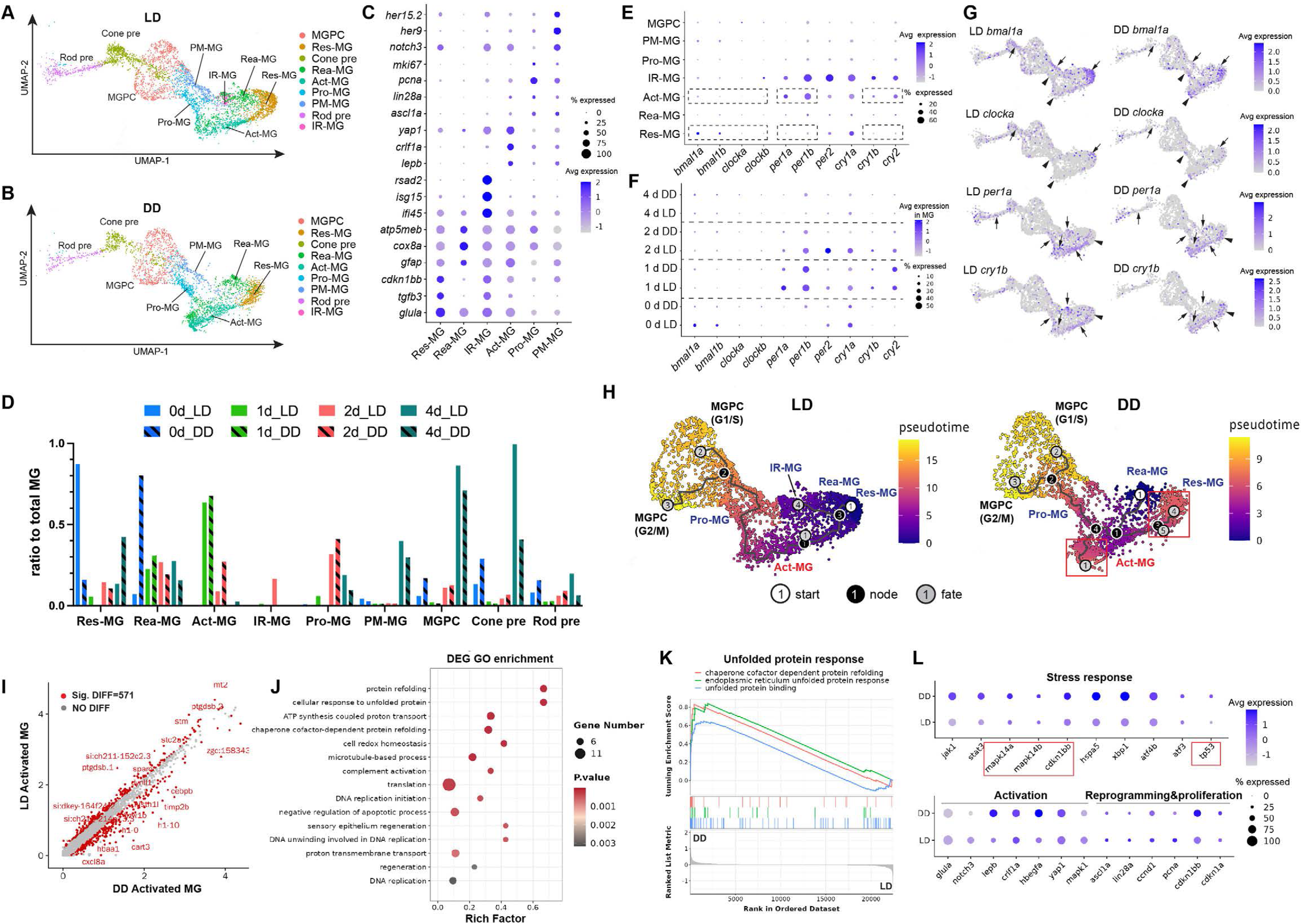
scRNA-seq analysis of MG reprogramming under LD or DD conditions. (A,B) UMAP visualization showing MG subtypes, MGPC, and photoreceptor precursors under LD or DD conditions. (C) Dot plot showing MG subtype-specific markers. (D) Ratio of MG subtypes, MGPCs and photoreceptor precursors to the total MG number at 0-4 dpi under LD or DD conditions. (E) Dot plot showing the expression of core clock genes in MG subtypes and MGPCs under LD conditions. (F) Dot plot showing the expression of core clock genes at different time points in MG under LD or DD conditions. (G) UMAP visualization showing the expression of core clock genes in MG, MGPCs and photoreceptor precursors under LD or DD conditions. Arrows and arrowheads in the LD panels indicate high and low expression, respectively. DD treatment disrupts clock gene expression at these locations. (H) Pseudotime analysis showing MG trajectories under LD or DD conditions. (I) Scatter plot showing differentially expressed genes (DEGs) in Act-MG under DD versus LD conditions. (J) GO enrichment of the DEGs in Act-MG. (K) GSEA of gene sets related to the unfolded protein response in Act-MG. (L) Dot plot showing the expression of genes related to stress response, MG activation, reprogramming, and proliferation under LD or DD conditions; Red boxes indicate cell cycle inhibitors. Res-MG, Resting MG; Rea-MG, Reactive MG; Act-MG, Activated MG; IR-MG, Immunoregulatory MG; Pro-MG, Proliferating MG; PM-MG, Post-mitotic MG. MGPC, MG-derived progenitors; Rod pre, Rod precursors; Cone pre, Cone precursors.

Under LD condition, core clock genes *bmal1a and bmal1b* were expressed in Res-MG in uninjured retinas, downregulated in Rea/Act/Pro-MG, and partially recovered in PM-MG in injured retinas (Figure 5E-G). *clocka* and *clockb* exhibited a similar expression pattern to *bmal1a* and *bmal1b*, albeit at lower levels (Figure 5E-G). In contrast, negative regulators *per1a*, *per1b*, *cry1b* and *cry2* showed complementary upregulation in Act/IR-MG (Figure 5E-G), suggesting that the core clock function was suppressed during MG reprogramming and proliferation. Notably, *per2* and *cry1a* exhibited an opposite expression pattern compared to other negative regulators (Figure 5E), consistent with previous reports showing that they were directly light-inducible in zebrafish^37^. Unlike negative regulators, *bmal1a, bmal1b*, *clocka and clockb* were expressed in some MGPCs (Figure 5E-G), suggesting clock involvement in MGPC proliferation or differentiation. Importantly, DD treatment disrupted core clock gene expression in MG at different timepoints (Figure 5F) and across MG subtypes (Figure 5G), further confirming the effectiveness of this method.

We next examined the impact of clock disruption on MG subtype composition and reprogramming trajectories. Uninjured LD retinas were dominated by Res-MG, whereas DD retinas showed mostly Rea-MG (Figure 5D, blue bars), suggesting clock disruption affected MG status before retinal injury. At 1 dpi, both LD and DD retinas consisted mainly of Rea/Act-MG, while a small proportion of LD MG had entered the cell cycle (Figure 5D, green bars). At 2 dpi, a higher percentage of DD MG remained in the Act-MG subgroup compared to LD, and the proportion of Pro-MG was similar in both groups (Figure 5D, red bars). Interestingly, the Immunoregulatory MG was only observed in 2 d LD retinas (Figure 5D, red bar). At 4 dpi, DD retinas showed fewer Pro/PM-MG, MGPCs and photoreceptor precursors, with a much higher proportion of MG returning into Res-MG (Figure 5D, cyan bars). Pseudotime analysis revealed two major LD trajectories (Res→Rea→IR; Res→Act→Pro→MGPC), whereas DD trajectories started from Rea-MG, with most cells returning to Res-MG or stalled in Act-MG, and a small proportion entering the cell cycle (Figure 5H).

As clock disruption significantly altered the Act-MG trajectory, we performed GO enrichment analysis on the 571 differentially expressed genes (DEGs) (Figure 5I) and GSEA between LD and DD Act-MG. GO analysis indicated significant difference in unfolded protein response (UPR), ATP/protein synthesis, cell redox homeostasis, DNA replication and regeneration pathways (Figure 5J). GSEA revealed upregulation of UPR-related gene sets in DD Act-MG (Figure 5K). These cells also exhibited upregulation of stress genes (including cell-cycle inhibitors *mapk14a/14b*, *cdkn1bb*, *tp53*) and MG activation genes, along with downregulation of reprogramming/proliferation genes (Figure 5L). GSEA further revealed downregulation of ATP/protein synthesis and DNA replication gene sets in DD Act-MG (Figure S3A-C), indicating a lack of growth signals. Consistently, p-S6 staining showed reduced mTORC1 activity—a master regulator of protein translation and metabolism—in DD MG (Figure S3D,E). Together, these findings demonstrated that clock disruption impaired MG reprogramming and induced stress-metabolic signatures.

### 3.6 Cell-cell communication analysis identifies the Cxcl12-Cxcr4 axis as a key chemotactic signaling between MG, vascular endothelial cells (EC) and immune cells

As clock disruption suppressed the innate immune responses in the retina, we examined microglia—key regulators of retinal regeneration—in LD and DD retinas. Clustering analysis identified two microglial subtypes: Micro-1 and Micro-2 (Figure 6A). Micro-1 showed higher expression of pro-inflammatory cytokines IL-1β (*il1b*) and TNF-α (*tnfb*), as well as Syndecan-4 (*sdc4*), a pro-inflammatory proteoglycan (Figure 6B), indicating a pro-inflammatory subgroup. In contrast, Micro-2 was almost completely absent in DD-retinas and only present at 2 dpi LD retinas (Figure 6A). This subtype expressed high levels of *pcna*, *igf2b*, *irf7* and *marco* (Figure 6B), with GO enrichment for proteasome, immune response and lysosome (Figure 6C), suggesting that it’s a proliferative subtype with immune-regulatory and reparative functions. Cell composition analysis revealed that DD treatment markedly reduced Micro-1 at 1 dpi and abolished Micro-2 at 2 dpi (Figure 6D), consistent with the above results (Figure 4). Importantly, both microglia subgroups expressed cytokines (*il1b*, *tnfb*, *il6*) and growth factors (*grn1/2*, *hbegfa/b*, *pdgfbab*) implicated in retinal regeneration^14,34,38–41^ (Figure 6E). Thus, reduced microglia accumulation and the subsequent loss of regeneration-associated secreted factors may underlie the regenerative defects in DD-treated retinas.

**Figure 6.**
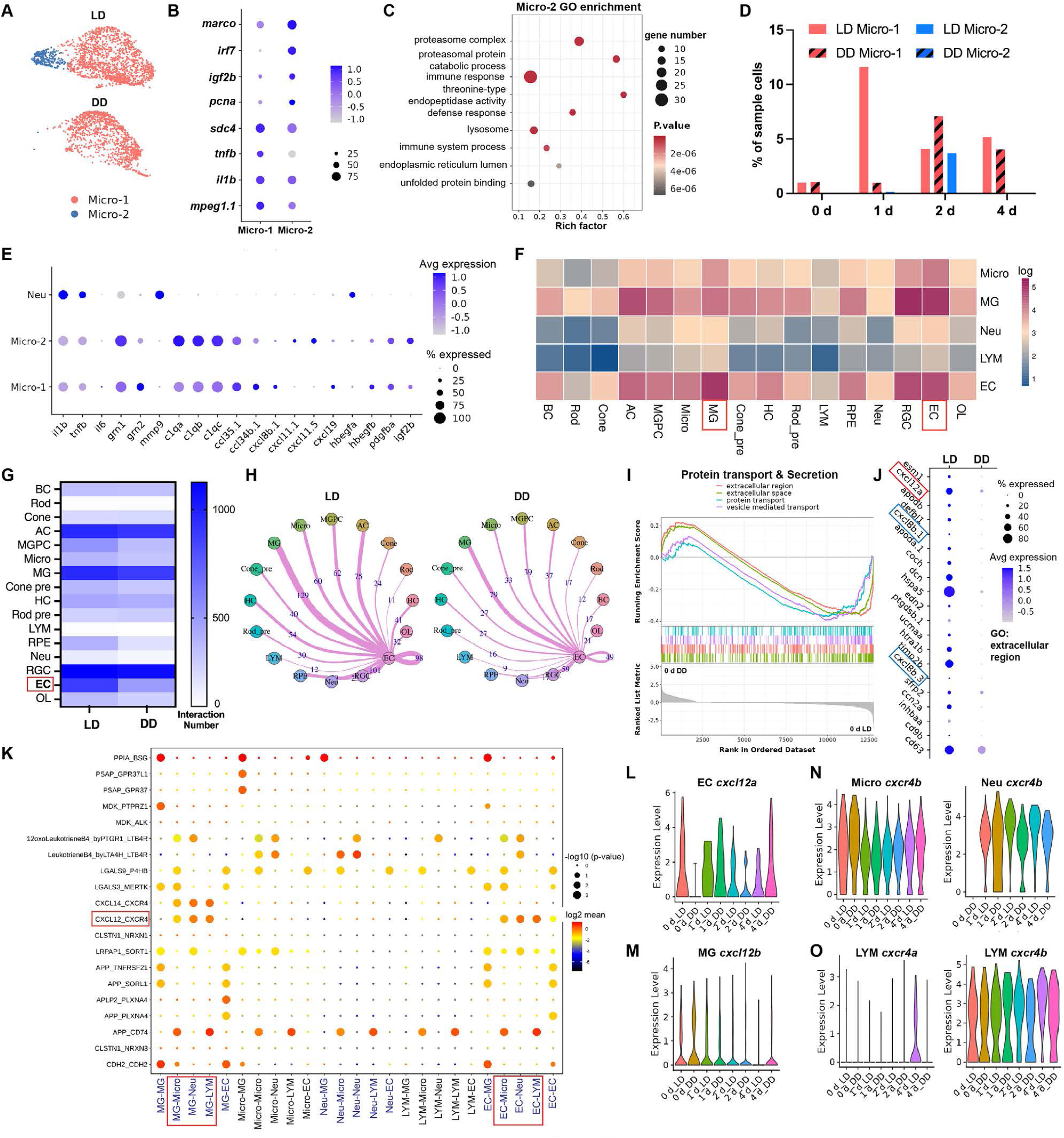
Cell-cell communication analysis identifies the Cxcl12-Cxcr4 chemotactic axis in the retina. (A,B) Microglia subtypes (Micro-1, Micro-2) and their specific markers. (C) GO enrichment of Micro-2 compared to Micro-1. (D) Proportion of microglia subtypes at 0-4 dpi under LD or DD conditions. (E) Dot plot showing the expression of secreted factors in microglia subtypes. (F) Predicted cell-cell interactions of MG, EC, and immune cells with all retinal cell types in LD retinas. (G) Total interaction numbers for each retinal cell type under LD or DD conditions. (H) Interaction of EC with other retinal cell types under LD or DD conditions. (I) GSEA of gene sets related to protein-transport and secretion in EC under LD or DD conditions. (J) Top 20 secreted factors significantly decreased in DD-treated EC compared to LD conditions. (K) Most significant ligand-receptors pairs between MG, EC, and immune cells. (L-O) Violin plots showing the expression of *cxcl12a/12b* and *cxcr4a/4b* in MG, EC, and immune cells at 0-4 dpi under LD or DD conditions. AC, amacrine cells; BC, bipolar cell; Cone, cone photoreceptor; Cone pre: cone precursors; EC, endothelial cell; HC, horizontal cell; LYM, lymphocyte; MG, Müller glia; MGPC, MG-derived progenitors; Micro, microglia; Neu, neutrophil; OL, oligodendrocyte; RGC, retinal ganglion cell; Rod, rod photoreceptor; Rod pre, rod precursors; RPE, retinal pigment epithelium.

To investigate immune regulation of MG reprogramming, we performed the CellphoneDB analysis of immune-MG crosstalk in LD and DD retinas. In LD retinas, the top three cell types interacting with MG are RGC, vascular endothelial cell (EC) and amacrine cells (Figure 6F, 2^nd^ row). ECs and MG are also among the top three partners of immune cells (Figure 6F). Given that peripheral immune cells (e.g., neutrophils, lymphocytes) must cross the blood-retina-barrier formed by ECs, we hypothesized that EC mediate MG-immune communication. Indeed, DD-treatment substantially reduced EC crosstalk with nearly all other retinal cell types (Figure 6G,H), accompanied by decreased protein transport and secretion (Figure 6I). Among the top 20 secreted factors differentially expressed by EC between LD and DD, we identified the chemokine Cxcl12 (Sdf-1, Figure 6J), a known potent leukocyte chemoattractant^42–44^. Furthermore, Cxcl12-Cxcr4 was among the top 20 significant ligand-receptor pairs connecting MG, EC and immune cells (Figure 5K). scRNA-seq showed that zebrafish Cxcl12 orthologues *cxcl12a* and *cxcl12b* were expressed by EC and MG, respectively, while their receptors *cxcr4a* and *cxcr4b* were dominantly expressed by immune cells (Figure 6L-O, Figure S4). Importantly, DD-treatment markedly downregulated *cxcl12a* expression in ECs in both uninjured and injured retinas (Figure 6L), suggesting that it was the dominant form responsible for the attenuated immune responses in clock-disrupted retinas.

### 3.7 Circadian clock-controlled Cxcl12 expression regulates the immune-and regenerative responses in the injured retina

To determine whether Cxcl12a/12b expression was under circadian clock control, we first assessed their mRNA expression in LD, DD or DL retinas. qPCR revealed rhythmic expression of both transcripts under LD conditions during the 0-48 hpi period (Figure 7A,B, zero-amplitude F-test, *p* < 0.05). Notably, both *cxcl12a* and *cxcl12b* peaked at ZT4, aligning with the pattern of retinal *bmal1a* and *clocka* expression. Importantly, both DD and DL treatment significantly reduced the amplitude (*p* < 0.01) and mesor (*p* < 0.01) of *cxcl12a* (Figure 7A). For *cxcl12b*, circadian disruption significantly reduced its amplitude (*p* < 0.05) and caused a phase advance (*p* < 0.05) (Figure 7B). These findings indicated that retinal *cxcl12a/12b* expression was under circadian control. To explore the underlying mechanism, we examined their promoter regions and identified two E-box elements upstream of the transcription start site (Figure 7C,E). Luciferase assays demonstrated that Bmal1b and Clocka significantly increased the promoter activity of both *cxcl12a* and *cxcl12b*, but Per2 repressed it (Figure 7C,E). Importantly, deletion of these E-box sequences abolished the transactivation of *cxcl12a/12b* promoters (Figure 7C,E). Moreover, chromatin immunoprecipitation (ChIP) assays showed that Bmal1b bound to the predicted E-box elements in *cxcl12a*/*12b* promoters (Figure 7D,F). These findings demonstrated that retinal *cxcl12a/12b* expression was directly controlled by the circadian clock via E-boxes.

**Figure 7.**
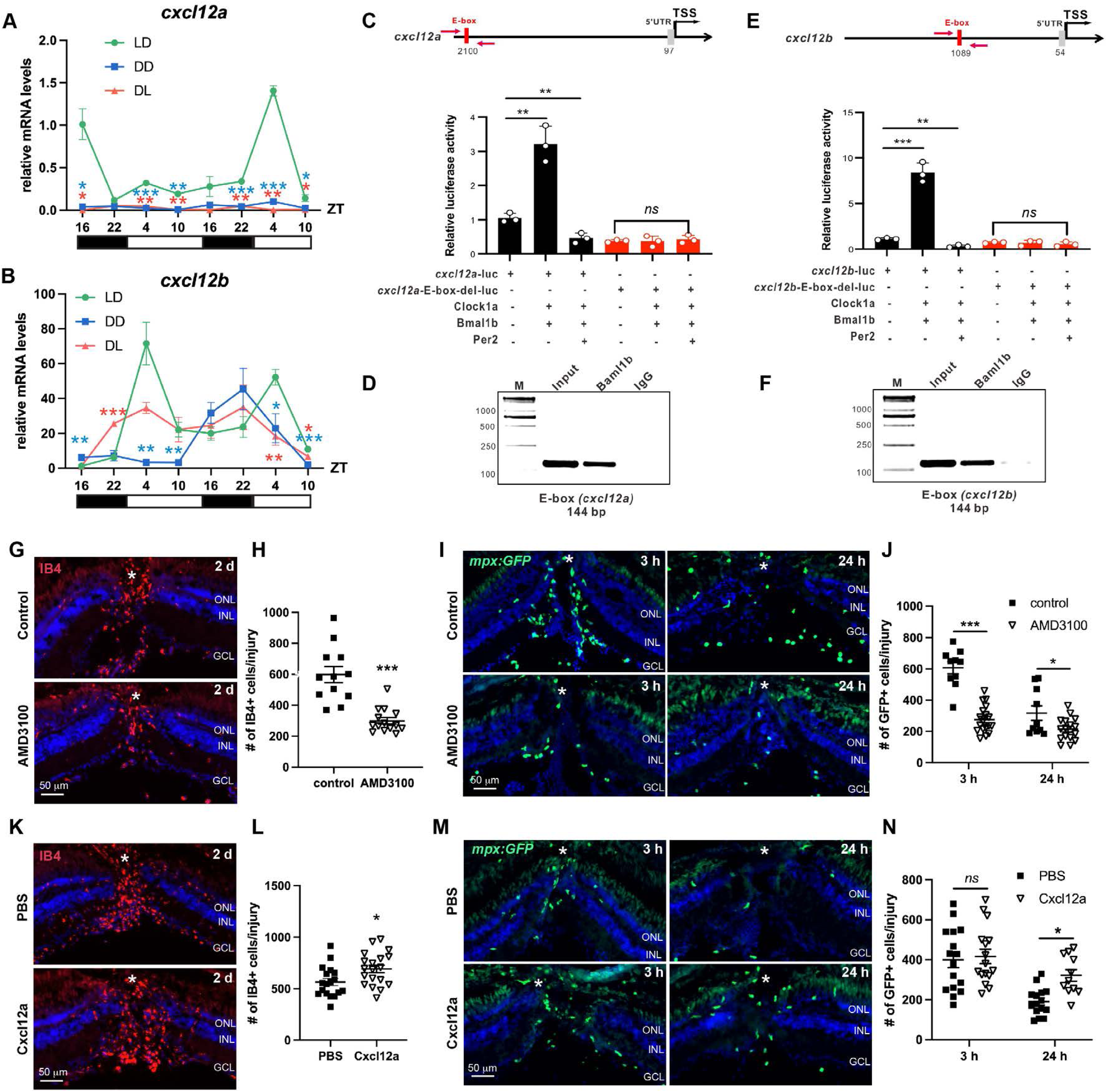
Clock-controlled Cxcl12 regulates innate immune responses in the injured retina. (A,B) qPCR analysis showing the expression of *cxcl12a* and *cxcl12b* from 0 to 48 hpi under LD, DD or DL conditions. Statistical differences were assessed by comparing DD or DL values to LD controls. (C,E) Schematic diagrams of the cxcl12a and cxcl12bs promoters containing the E-box element (red). Luciferase assays showing the regulation of WT or E-box-mutant *cxcl12a/12b* promoter activities by Bmal1b, Clocka, or Per2. *n* = 3. (D,F) ChIP assays targeting the E-box elements in the *cxcl12a/12b* promoters. (G,H) IB4 staining of microglia at the injury site in control or AMD3100-treated retinas (G) and quantification (H). (I,J) Distribution of neutrophils in control or AMD3100-treated retinas of the *Tg(mpx:GFP)* transgenic line (I) and quantification (J). (K,L) IB4 staining of microglia in PBS-or Cxcl12a-injected retinas (K) and quantification (L). (M,N) Neutrophil accumulation at the injury site in PBS- or Cxcl12a-injected retinas (M) and quantification (N). White asterisks mark the needle-poke injury sites. *ns*, not significant; *, *p* < 0.05; **, *p* < 0.01; ***, *p* <0.001. TSS: transcription start site; ONL, outer nuclear layer; INL, inner nuclear layers; GCL, ganglion cell layer.

We next investigated whether the Cxcl12-Cxcr4 signaling modulated the immune response following retinal injury. Indeed, treatment with a Cxcr4 antagonist AMD3100 significantly reduced the number of microglia at the injury site at 2 dpi (Figure 7G,H). AMD3100 also reduced neutrophil number at the injury site during both the peak (3 hpi) and resolution (24 hpi) phases (Figure 7I,J). Conversely, intravitreal injection of recombinant zebrafish Cxcl12a significantly increased the number of microglia in the injured region (Figure 7K-N). Cxcl12a injection did not affect neutrophil numbers at 3 hpi but increased them at 24 hpi (Figure 7M,N), suggesting that Cxcl12-Cxcr4 signaling suppressed neutrophil resolution in injured zebrafish retinas.

To examine whether the circadian clock regulated the retinal immune and regenerative responses via Cxcl12, DD retinas received intravitreal Cxcl12a injection at the time of injury, with PBS-injected DD or LD retinas as controls. Cxcl12a injection significantly increased microglia numbers in injured retinas compared to DD controls, reaching levels comparable to LD controls (Figure 8A,B). Cxcl12a injection induced a slight, non-significant increase in neutrophil numbers at 6 hpi (Figure 8C,D). qPCR revealed that Cxcl12a injection restored the expression of *il1b* and *il6*, but not *il11a*, *tnfa*, or *tnfb*, in DD retinas (Figure 8E, upper panel), indicating partial rescue of the inflammatory response. Furthermore, Cxcl12a injection restored retinal expression of key MG reprogramming factors *ascl1a*, *lin28a* and *lepb* in DD retinas (Figure 8E, lower panel). Importantly, Cxcl12a injection significantly increased MGPC numbers compared to DD controls, though levels remained below those in LD retinas (Figure 8F,G). Together, these results demonstrated that Cxcl12a injection partially rescued the defective immune and regenerative responses in DD-treated retinas, indicating that the circadian clock regulated these responses in a Cxcl12-dependent manner.

**Figure 8.**
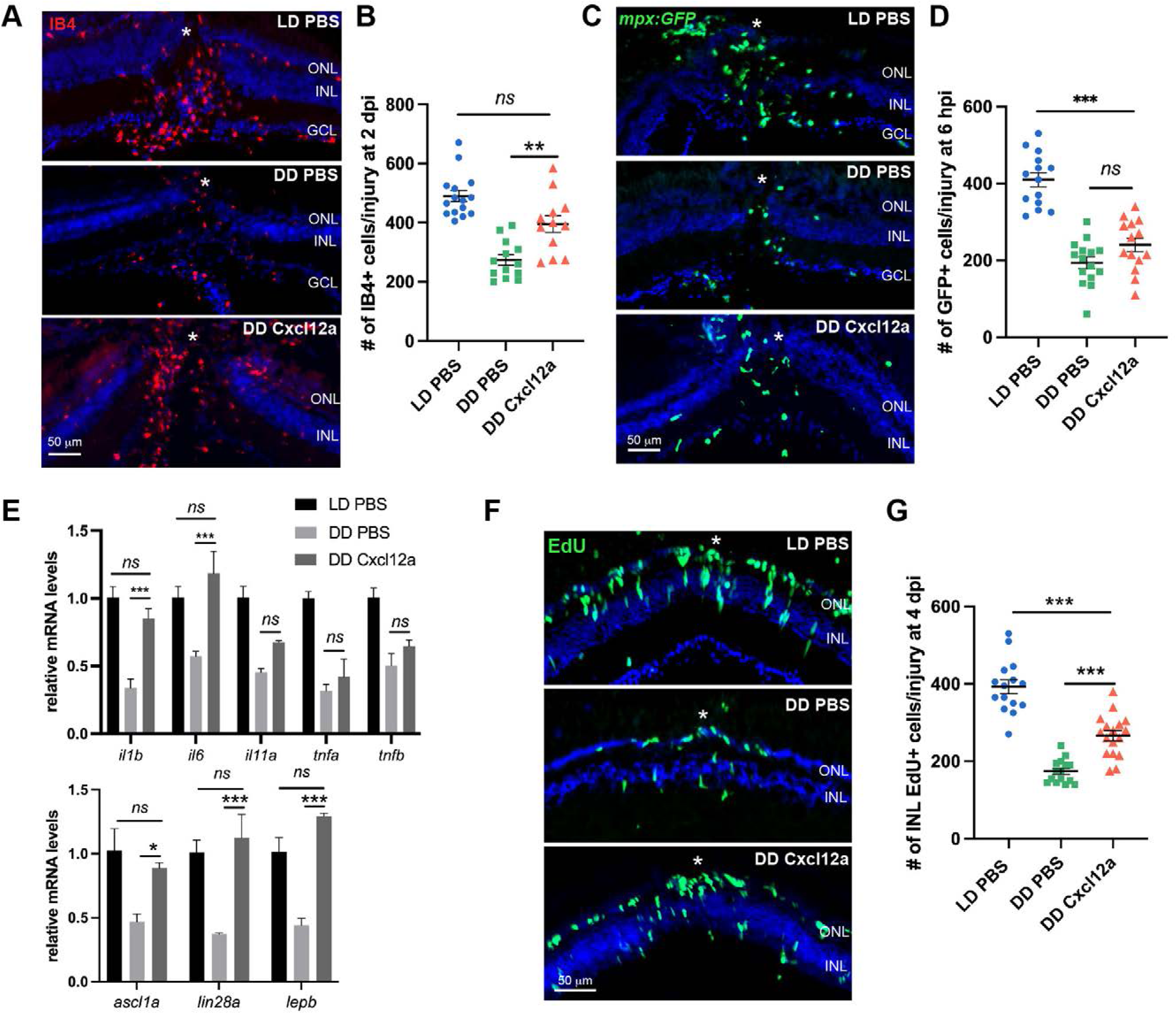
Cxcl12a injection partially restores immune-regenerative responses in DD-treated retinas. (A,B) IB4 staining of retinal microglia in the LD PBS, DD PBS or DD Cxcl12a groups at 2 dpi (A) and quantification (B). (C,D) Distribution of neutrophils in the retina of the *Tg(mpx:GFP)* transgenic line in the indicated groups at 6 hpi (C) and quantification (D). (E) qPCR analysis showing the expression of inflammatory cytokines (upper panel) and MG reprogramming genes (lower panel) in retinas at 2 dpi. (F,G) EdU immunostaining showing MGPCs in the retina at 4 dpi (F) and quantification (G). White asterisks mark the needle-poke injury sites. *ns*, not significant; *, *p* < 0.05; **, *p* < 0.01; ***, *p* <0.001. ONL, outer nuclear layer; INL, inner nuclear layers; GCL, ganglion cell layer.

## 4 DISCUSSION

In this study, using the zebrafish retinal regeneration model, we investigated the role of circadian clock in vertebrate CNS repair. We observed that regenerative responses differed when injury occurred during the day versus night, and showed that both regenerative and inflammatory responses exhibited diurnal fluctuations, peaking during the rest phase (ZT22). Importantly, disrupting the clock via light manipulation markedly suppressed immune and regenerative activity, as well as subsequent neuronal regeneration. scRNA-seq revealed aberrant MG reprogramming characterized by stress and metabolic signatures. Cell-cell communication analysis identified Cxcl12-Cxcr4 signaling as a key interaction between immune cells, EC and MG. Mechanistically, retinal Cxcl12 expression was under circadian control, with core clock components directly binding E-box elements in the *cxcl12a/12b* promoters to activate transcription. Functionally, the circadian clock regulated immune and regenerative responses in a Cxcl12-dependent manner. Collectively, this study uncovers a novel circadian role in vertebrate CNS repair, with potential implications for regenerative therapies in the mammalian CNS.

The circadian clock is well-known for its important role in DNA synthesis and cell-cycle progression^45^. Previous studies have shown that the circadian clock regulates neural stem cell (NSC) proliferation and neurogenesis^46,47^, and its dysregulation is associated with neurodegenerative diseases such as sleep disorder, anxiety and mood alteration^48,49^. Similar to NSC, MG function as retinal stem cells in the injured zebrafish retina^50^, undergoing asymmetric division to generate MGPCs while self-renewing^51^. Consistently, our data show that MG/MGPC proliferation exhibits rhythmic oscillations, and clock disruption significantly suppresses their proliferation. Furthermore, scRNA-seq analysis reveals that DD treatment disrupts MG reprogramming trajectories, causing a large proportion of Act-MG to either stall in this state or return to Res-MG. At the molecular level, clock disruption leads to increased unfolded protein response (UPR) and stress signaling, concomitant with reduced ATP production, protein synthesis and DNA replication in Act-MG. Notably, in the LD retinas, the core clock genes *bmal1a/1b* and *clocka/b* are downregulated in Act-MG, whereas the negative regulators *per1a/1b* and *cry1b/2* are markedly upregulated (Figure 5E,G), indicating suppressed clock function during MG reprogramming. Given that stress and inflammation are known to disrupt normal clock function^52–55^, these findings suggest that injury signals rewire the circadian machinery from a homeostatic oscillatory mode to a stress-activated state.

The circadian clock plays a pivotal role in immune system regulation^18,19^. Numerous studies have shown that the clock influences many aspects of immune cell function—including cell proliferation, migration and phagocytosis— by regulating cellular metabolism, chemokine/chemokine receptor expression, and cytokine production^56^. Consistently, our results indicate that circadian disruption compromises retinal immune responses, and that enhancing inflammation via zymosan injection restores the regenerative response (Figure 4), underscoring the pivotal role of clock-regulated immunity in retinal regeneration. The circadian clock is known to regulate the expression of many chemokines, such as CCL2 in muscle stem cell-neutrophil crosstalk^57^, CXCL14 in skin innate immunity^58^, CCL2/CCL8 and S100A8 in monocyte recruitment^59^, CXCL2 in neutrophil aging^60^, and CXCL5 in pulmonary neutrophil recruitment^61^. The clock also regulates CXCL12-controlled cyclic release of hematopoietic stem cells into the bloodstream through circadian noradrenaline secretion by the sympathetic nervous system^62^. However, whether the clock regulates tissue regeneration in Cxcl12-dependent immune modulation remains unclear. In this study, we report that retinal *cxcl12a/12b* expression is under circadian control, and that Bmal1 directly activates their transcription via binding to the E-box elements in their promoters (Figure 7A-F). This conclusion is supported by the near-synchronous expression of *cxcl12a/12b, bmal1a,* and *clocka* in the retina, which peaked at ZT4 and reached a nadir around ZT22 (Figure 7A,B). Compared with *cxcl12a/12b* mRNA oscillations, microglia accumulation and inflammatory cytokine expression in the injured retina appear to peak later at around ZT22 (Figure 1F,4B). This delay is likely attributable to the time required for Cxcl12 protein translation and secretion, Cxcr4 binding and downstream signaling, and ultimately microglia activation and recruitment. Our scRNA-seq revealed that EC and MG are the primary source of retinal *cxcl12a* and *cxcl12*b, respectively (Figure S4). ECs are known to express CXCL12, which is crucial for angiogenesis during embryonic development, wound repair and pathological conditions^63–65^. However, CXCL12 expression by MG has not been reported. Therefore, future studies should further investigate the expression patterns and specific functions of these two Cxcl12 homologs in retinal regeneration.

The chemokine CXCL12 is a potent chemoattractant for leukocytes in both physiological and pathological conditions^42,66^. The CXCL12-CXCR4 signaling has been shown to promote the chemotaxis and recruitment of microglia^67^, macrophage^44^ and monocyte^68–70^. Consistently, we report that the CXCR4 antagonist AMD3100 significantly reduced microglia recruitment in the injured retina, while intravitreal injection of zebrafish Cxcl12a increased their number (Figure 6G,H,K,L). Our findings support a pivotal role for Cxcl12 in microglia recruitment during retinal regeneration. The effects of CXCL12 on neutrophil chemotaxis are more complex, as previous studies suggest that it can retain neutrophils in the bone marrow and inflammatory site^71,72^, and also promote their recruitment to certain tissues^70,73^. In this study, we investigated the role of Cxcl12-Cxcr4 in neutrophil recruitment/retention in the injured retina at two timepoints: the peak of neutrophil recruitment at 3 hpi, and the resolution phase at 24 hpi. We show that local AMD3100 treatment significantly decreased the neutrophil number at both timepoints, while Cxcl12a injection increased their number at 24 hpi but had no effect at 3 hpi (Figure 6I,J,M,N). If Cxcl12a is a recruitment signal, then its injection would increase neutrophil number at both timepoints. It’s therefore more likely that Cxcl12a functions as a retaining signal for neutrophils at the injury site, rather than a direct recruitment signal. Nevertheless, as there is only one commercially available zebrafish Cxcl12 protein (Cxcl12a) and both homologs can signal through Cxcr4b^74,75^, it’s also possible that Cxcl12b alone or a combination of both may have different results and thus require further investigation.

Several limitations warrant consideration. First, the lack of MG-specific clock mutants prevents definitive conclusions about cell-autonomous vs non-cell-autonomous clock functions in MG reprogramming. Second, the absence of reliable zebrafish Cxcl12a/12b antibodies limits protein-level validation. Third, Cxcl12a injection in DD retinas achieved only partial rescue of immune and regenerative responses, suggesting additional clock-regulated factors (e.g., Cxcr4 expression on immune cells^76^, other clock-controlled chemokines) contribute to these responses. Fourth, whether circadian gating of regeneration applies to other CNS regions or mammalian systems remains to be tested.

In summary, we establish that the circadian clock gates CNS regeneration by orchestrating Cxcl12-dependent immune coordination. This circadian-immune axis represents a previously unrecognized regulatory node in retinal repair. Our findings suggest that chronotherapeutic strategies—timing interventions to align with circadian rhythms—could enhance CNS repair in mammals.

## ACKNOWLEDGEMENT

This study was supported by grants from the National Natural Science Foundation of China (81970820, 82371062, 32571347), and Basic Research Program of Jiangsu Education Department (24KJA180008).

## DATA AVAILABILITY STATEMENT

The data that support the findings of this study are available on request from the corresponding author.

