## Supplemental Material for "Circadian clock gates retinal regeneration by orchestrating Cxcl12-dependent immune coordination"

#### Table 1. Primers used for qPCR.

| Gene name | Forward primer (5’-3’) | Reverse Primer (5’-3’) |
|---|---|---|
| *ascl1a* | CAGAACAGGTCGAGAGTCCG | TCATCTTCTTGTTGGCCGCT |
| *bmal1a* | GACATCATGGAGGAGCCTGG | TCCACTGCTGATACGTTTACACT |
| *clocka* | CAGGCCCCACGTCTTTTACA | TTTACTGAGGCGGAGGGTTG |
| *ccnd1* | CTGGACAGGTTTTTATCTGTGGAGCC | GCTTGGAGCTCTGATGTATAGGCAGT |
| *ccne1* | CACGTTAAGGCTCTCGACATTCAAG | GCATGGGCTTGTGTAACCTGTGT |
| *cdk1* | CTGGCAGATTTCGGCTTAGCCCGTGC | CTTATAGTCTGGCAGAGACTCAACATCTGGC |
| *cdk2* | CTTAAACCCCAGAATCTCCTCATCAA | CTTAAACCCCAGAATCTCCTCATCAA |
| *cry1a* | CATTGGCAGCACTCACATGC | TAGTCTCCGTTGGGGTCAGT |
| *crlf1a* | GGGATTCTGGGATCTAGGAAAGC | TCCTTGAAGAACCTGGTTGCG |
| *cxcl12a* | CATGCACCGATTTCCAACGG | TTGGCACTGGAAGGGAGAAG |
| *cxcl12b* | CCCAGAGACTGACGGTAGGA | AGCTTGGCACTGGTAATGAGA |
| *hbegfa* | ATGTCTGACCATCATTGGCCTCC | ACCATTCAGCTTGCTGTGCCC |
| *il1b* | CTGGAGATGTGGACTTCGCA | AGTGCAGTCCTGGTAGTAGGT |
| *il6* | GCTATTCCTGTCTGCTACACTGG | TGAGGAGAGGAGTGCTGATCC |
| *il11a* | CTCCTCATCGCTGCTTCTCTCG | TTGCGAAGTCACTGGCTCTGC |
| *il11b* | ATTCGCTATCATCCCTGCCC | TAGGTGACAGACAGCACAGTTG |
| *lepb* | CATTGCTCGAACCACCATCAGC | TCTTTATGCACCGGGGTCTCG |
| *lin28a* | GGATGGGCTTCGGATTTCTGTC | TCCTCCACAGTTGAAGCATCGATC |
| *pcna* | TTACATTCAGAGCCTCGCCA | CCAGACGTGCCTCAAACATT |
| *per1b* | GCCATGCAGAAACATCAGCC | AGGCACCCATTTCTTCCTCG |
| *rpl13* | TCTGGAGGACTGTAAGAGGTATGC | AGACGCACAATCTTGAGAGCAG |
| *tnfa* | GGTGAAAATCCTGCCCCGTA | AATGGATGGCAGCCTTGGAA |
| *tnfb* | CCTCTCTATTCACAGGTGACCC | TGGAATGCCTGATCCACACC |

#### Table 2. Primers used for the luciferase assay and ChIP.

| name | Forward primer (5’-3’) | Reverse Primer (5’-3’) |
|---|---|---|
| *cxcl12a-luc* | CTGGCCTAACTGGCCGGTACCGT TTTCTTAACACGAATCATTCAAA | CTTGATATCCTCGAGGCTAGCCGTT GGAAATCGGTGCATGAATGGC |
| *cxcl12b-luc-f* | CTGGCCTAACTGGCCGGTACCAT GCAGCAGAAATTACTGACTCTAA | CTTGATATCCTCGAGGCTAGCTTGA TCACAGCGAGTGTCCAGAGCT |
| *cxcl12a-chip* | GTTTTCTTAACACGAATCAT | GATTCATTTGCTGGAATAGT |
| *cxcl12b-chip* | CTGGTGAGCACAAAATAAGA | GTTTAAATTCAGCCCATAGA |
| *cxcl12a-e-box-del* | TAACACGAATCATTCAAATCTCACTA TTAATTCGATTATT | AATAATCGAATTAATAGTGAGATTTGAA TGATTCGTGTTA |
| *cxcl12b-e-box-del* | CTATGGAAGTCCATGGCTTCTAAGG TAATTTTTTGCAAAA | TTTTGCAAAAAATTACCTTAGAAGCCA TGGACTTCCATAG |

#### Supplementary Figure Legends

**Figure S1. DL treatment suppressed the innate immune responses in injured retinas**. (A) IB4 staining of retinal microglia at 1 and 2 dpi under LD or DL conditions. (B) Quantification of microglia at the injury site in (A). (C) Distribution of neutrophils in retinas at 3-6 hpi from the *Tg(mpx:GFP)* transgenic line under LD or DL conditions. (D) Quantification of neutrophil in (C). White *, site of needle poke injury. ***, $p$ <0.001. ONL, outer nuclear layer; INL, inner nuclear layers; GCL, ganglion cell layer.

**Figure S2. scRNA-seq analysis of retinal cell types.** (A) tSNE showing the clustering of all retinal cell types in LD retinas. (B) Dotplot showing cell type specific markers. AC, amacrine cells; BC, bipolar cell; Cone, cone photoreceptor; Cone pre: cone precursors; EC, endothelial cell; HC, horizontal cell; LYM, lymphocyte; MG, Müller glia; MGPC, MG-derived progenitors; Micro, microglia; Neu, neutrophil; OL, oligodendrocyte; RGC, retinal ganglion cell; Rod, rod photoreceptor; Rod pre, rod precursors; RPE, retinal pigment epithelium.

**Figure S3. Lack of growth signals in Act-MG under DD conditions.** (A-C) GSEA analysis of gene sets related to ATP/protein synthesis and DNA replication in Act-MG under LD or DD conditions; (D) p-S6 and EdU immunofluorescence showing the mTORC1 activity in MG from injured LD or DD retinas. (E) Quantification of the p-S6$^+$ MG in (D). White *, site of needle poke injury. **, $p$ < 0.01; ***, $p$ <0.001. ONL, outer nuclear layer; INL, inner nuclear layers; GCL, ganglion cell layer.

**Figure S4. scRNA-seq analysis of *cxcl12a/12b* and *cxcr4a/4b* expression in retinal cell types.** (A) Dotplot showing *cxcl12a/12b* and *cxcr4a/4b* expression in various cell types in LD retinas. (B) UMAP showing the expression of these genes in retinal cell types from LD retinas. AC, amacrine cells; BC, bipolar cell; Cone, cone photoreceptor; Cone pre: cone precursors; EC, endothelial cell; HC, horizontal cell; LYM, lymphocyte; MG, Müller glia; MGPC, MG-derived progenitors; Micro, microglia; Neu, neutrophil; OL, oligodendrocyte; RGC, retinal ganglion cell; Rod, rod photoreceptor; Rod pre, rod precursors; RPE, retinal pigment epithelium.

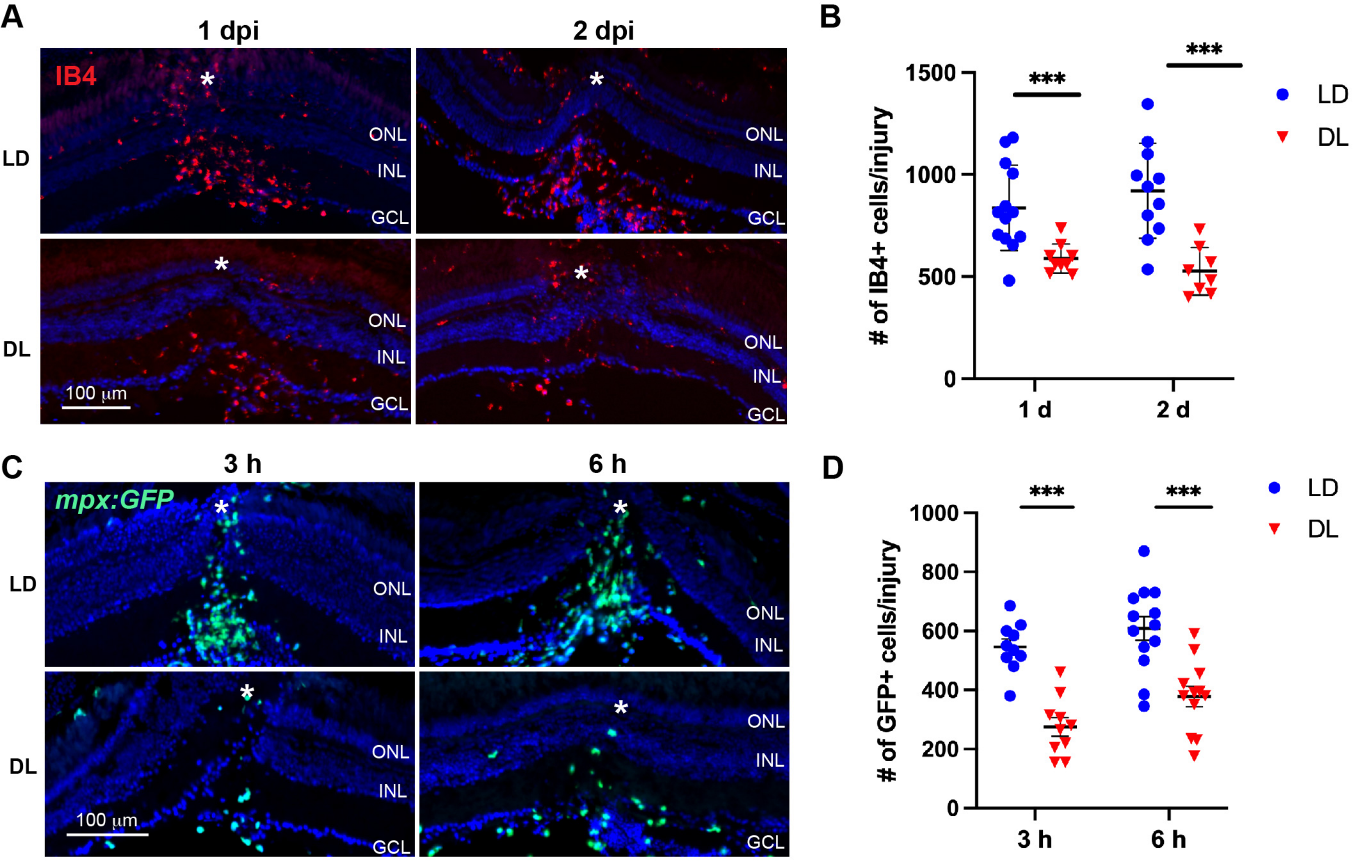
A
1 dpi
2 dpi
IB4
LD
DL
ONL
INL
GCL
100 μm
B
### of IB4+ cells/injury
1500
1000
500
0
1 d
2 d
***
LD
DL
C
3 h
6 h
mpx:GFP
D
### of GFP+ cells/injury
1000
800
600
400
200
0
3 h
6 h

**Figure S1**

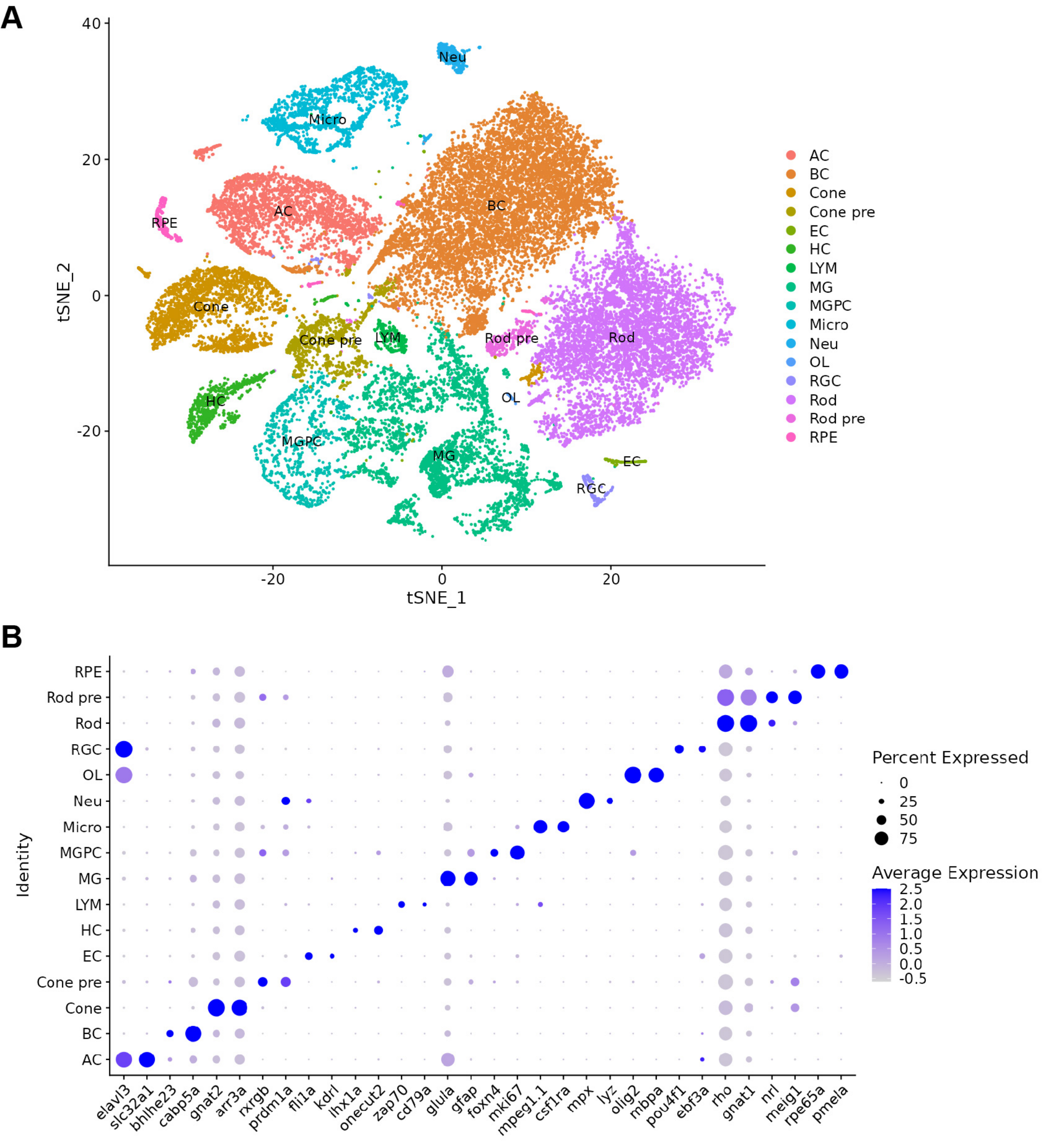
A
40
20
0
-20
tSNE_2
-20
0
20
tSNE_1
Neu
Micro
AC
BC
RPE
Cone
Cone pre
LYM
Rod pre
Rod
HC
OL
MGPC
MG
EC
RGC
AC
BC
Cone
Cone pre
EC
HC
LYM
MG
MGPC
Micro
Neu
OL
RGC
Rod
Rod pre
RPE
B
Identity
RPE
Rod pre
Rod
RGC
OL
Neu
Micro
MGPC
MG
LYM
HC
EC
Cone pre
Cone
BC
AC
elavl3
slc32a1
bhlhe23
cabp5a
gnat2
arr3a
rxrgb
prdm1a
fli1a
kdrl
lhx1a
onecut2
zap70
cd79a
glula
gfap
foxn4
mki67
mpeg1.1
csf1ra
mpx
lyz
olig2
mbpa
pou4f1
ebf3a
rho
gnat1
nrl
meig1
rpe65a
pmela
Percent Expressed
0
25
50
75
Average Expression
2.5
2.0
1.5
1.0
0.5
0.0
-0.5

### Figure S2

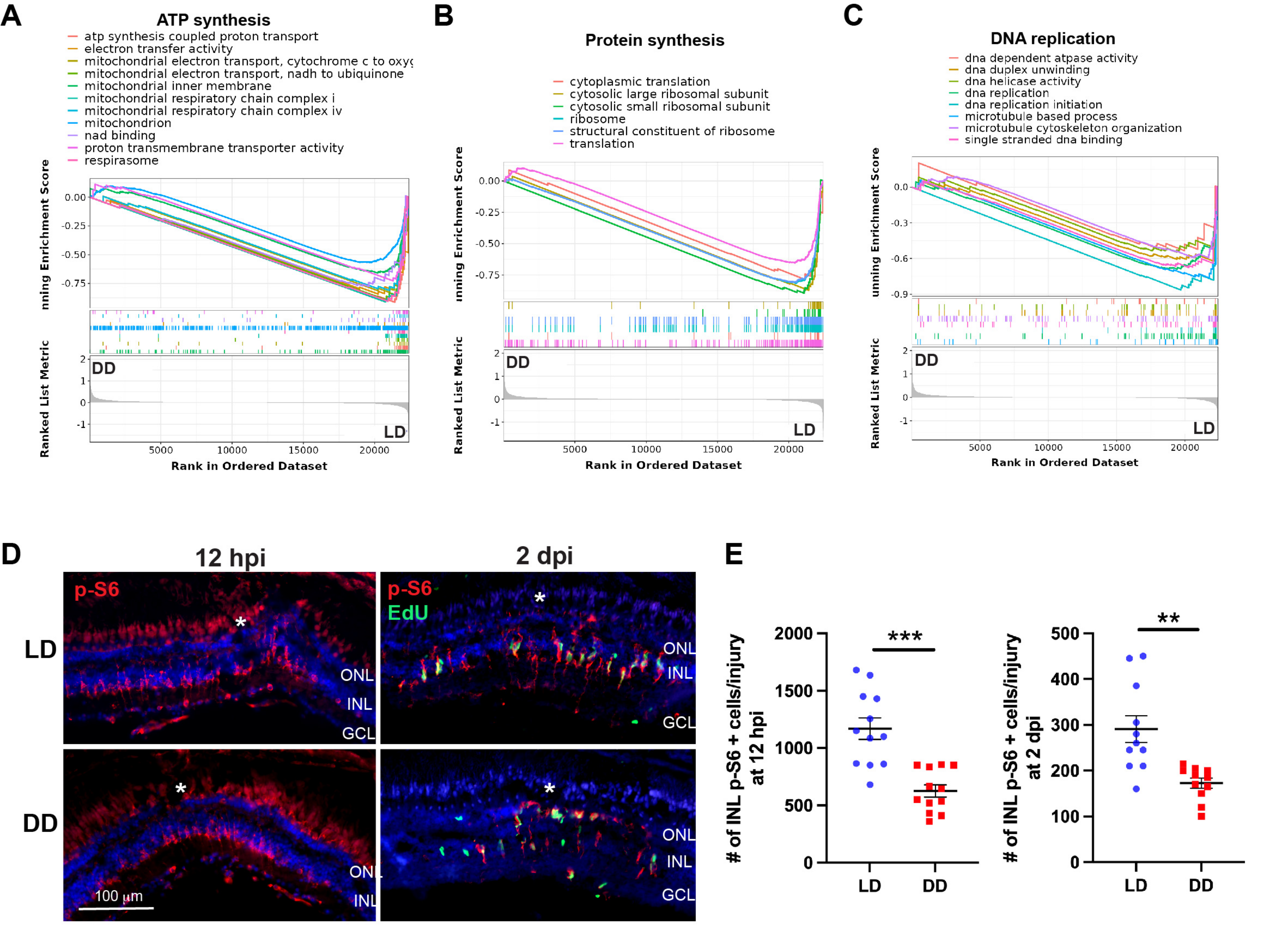
A
ATP synthesis
atp synthesis coupled proton transport
electron transfer activity
mitochondrial electron transport, cytochrome c to oxyg
mitochondrial electron transport, nadh to ubiquinone
mitochondrial inner membrane
mitochondrial respiratory chain complex i
mitochondrial respiratory chain complex iv
mitochondrion
nad binding
proton transmembrane transporter activity
respirasome
B
Protein synthesis
cytoplasmic translation
cytosolic large ribosomal subunit
cytosolic small ribosomal subunit
ribosome
structural constituent of ribosome
translation
C
DNA replication
dna dependent atpase activity
dna duplex unwinding
dna helicase activity
dna replication
dna replication initiation
microtubule based process
microtubule cytoskeleton organization
single stranded dna binding
Running Enrichment Score
Ranked List Metric
Rank in Ordered Dataset
DD
LD
D
12 hpi
2 dpi
p-S6
EdU
LD
DD
ONL
INL
GCL
100 µm
E
### of INL p-S6 + cells/injury at 12 hpi
### of INL p-S6 + cells/injury at 2 dpi
***
**

Figure S3

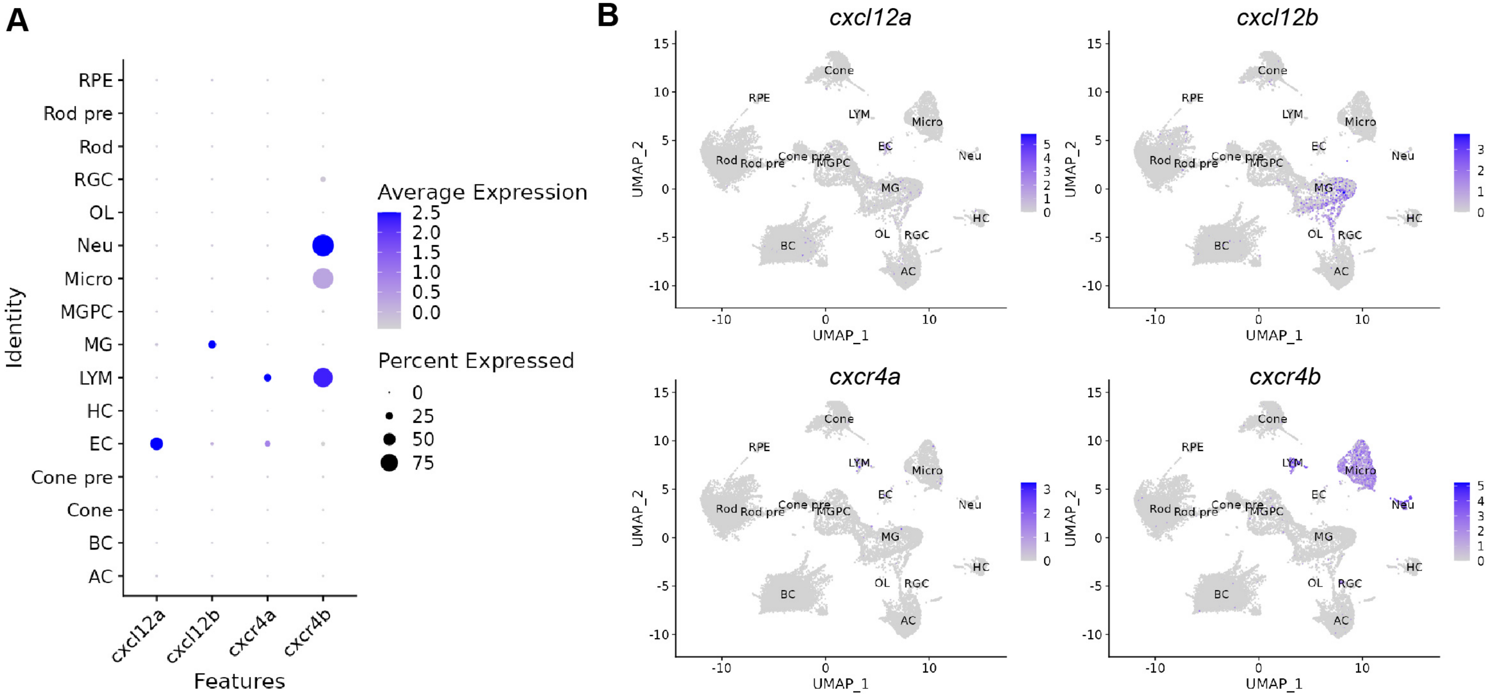
A
RPE
Rod pre
Rod
RGC
OL
Neu
Micro
MGPC
MG
LYM
HC
EC
Cone pre
Cone
BC
AC
Identity
cxcl12a
cxcl12b
cxcr4a
cxcr4b
Features
Average Expression
2.5
2.0
1.5
1.0
0.5
0.0
Percent Expressed
0
25
50
75
B
cxcl12a
cxcl12b
cxcr4a
cxcr4b
UMAP_1
UMAP_2
Cone
RPE
LYM
Micro
EC
Neu
Rod
Rod pre
Cone pre
MGPC
MG
HC
OL
RGC
BC
AC

**Figure S4**
